# Oligomerization of the FliF domains suggests a coordinated assembly of the bacterial flagellum MS ring

**DOI:** 10.1101/2021.09.23.461508

**Authors:** Giuseppina Mariano, Raquel Faba-Rodriguez, Soi Bui, Weilong Zhao, James Ross, Svetomir Tsokov, Julien Bergeron

## Abstract

The bacterial flagellum is a complex, self-assembling macromolecular machine that powers bacterial motility. It plays diverse roles in bacterial virulence, including aiding in colonization and dissemination during infection. The flagellum consists of a filamentous structure protruding from the cell, and the basal body, a large assembly that spans the cell envelope. The basal body is comprised of over 10 different proteins, forming several concentric ring structures, termed the M- S- L- P- and C-rings, respectively. In particular, the MS rings are formed by a single protein FliF, which consists of two trans-membrane helices anchoring it to the inner membrane and surrounding a large periplasmic domain. Assembly of the MS ring, through oligomerization of FliF, is one of the first steps of basal body assembly.

Previous computational analysis had shown that the periplasmic region of FliF consists of three structurally similar domains, termed Ring-Building Motif (RBM)1, RBM2 and RBM3. The structure of the MS-ring has been reported recently, and unexpectedly shown that these three domains adopt different symmetries, with RBM3 having a 34-mer stoichiometry, while RBM2 adopts two distinct positions in the complex, including a 23-mer ring. This observation raises some important question on the assembly of the MS ring, and the formation of this symmetry mis-match within a single protein. In this study, we analyze the oligomerization of the individual RBM domains in isolation, in the *Salmonella typhimurium* FliF orthologue. We demonstrate that the periplasmic domain of FliF assembles into the MS ring, in the absence of the trans-membrane helices. We also report that the RBM2 and RBM3 domains oligomerize into ring structures, but not RBM1. Intriguingly, we observe that a construct encompassing RBM1 and RBM2 is monomeric, suggesting that RBM1 interacts with RBM2, and inhibits its oligomerization. However, this inhibition is lifted by the addition of RBM3. Collectively, this data suggests a mechanism for the controlled assembly of the MS ring.

## 1 Introduction

The flagellum is a complex macromolecular motor, whose role is to allow swimming motility, through the rotation of a long filament at the bacterium cell surface. The flagellum is employed by many bacteria to swim in liquid environments (Minamino and Imada, 2015), but it also represents an important virulence factor, playing central roles in cell adhesion and invasion, secretion of other virulence factors and biofilm formation (Duan et al., 2013). The bacterial flagellum can be divided in four major regions: On the cytosolic side, the rotor and stators are responsible for inducing filament rotation, using the proton motor force or sodium gradient (depending on the bacterial species)(Berg, 2003; Li et al., 2011). The basal body is the region that spans the cell envelope, and includes consecutive ring-like structures, termed M-, S-, L-, and P-rings; the hook is a bended structure that protrudes from the basal body on the cell surface; and the filament is a long (up to several μm) tubular structure of > 10,000s copies of a singular protein, the flagellin (Nakamura and Minamino, 2019).

The M and S rings are formed by the protein FliF, a 60 kDa protein, embedded in the cytoplasmic membrane through two trans-membrane helices (Figure 1a). It oligomerizes into a circular membrane-spanning complex, forming the fundamental scaffold for flagellar structure and assembly (Minamino et al., 2008). In-between the two transmembrane helices, FliF possesses a large periplasmic region consisting of three globular domains termed Ring-Building Domains (RBM1, RBM2 and RBM3 respectively) (Figure 1a) (Bergeron, 2016). Those RBMs possess a common fold (Spreter et al., 2009), and possess structural homology with components of the Type III Secretion System (T3SS) injectosome, and in particular RBM1 and RBM2 have sequence similarity with the T3SS protein SctJ (Yip et al., 2005; Bergeron et al., 2015; Bergeron, 2016). Conversely, RBM3 shows homology to the SpoIIIAG protein (Bergeron, 2016; Zeytuni et al., 2017), a macromolecular complex involved in spore formation.

**Figure 1.**
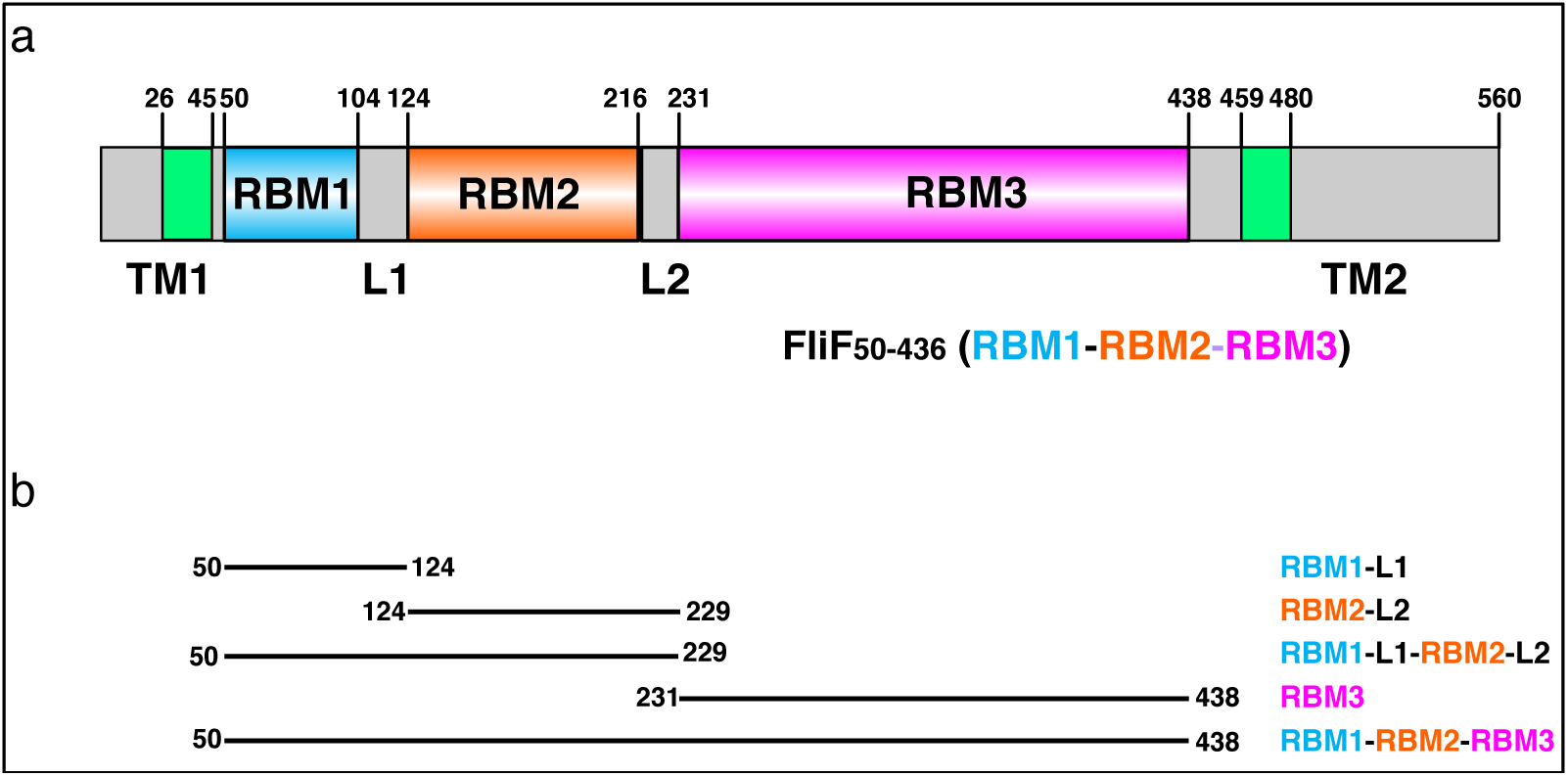
*Salmonella typhimurium* FliF domain organization. (a) Schematic representation of the domain composition and their boundaries in FliF. (b) Summary of the constructs, encompassing distinct FliF regions, used in this study.

On the cytosolic side, FliF binds to the protein FliG, part of the C-ring (Kubori et al., 1992; Levenson et al., 2012), via its C-terminus. This interaction is necessary for flagellum assembly (Li and Sourjik, 2011; Morimoto et al., 2014). FliG, together with FliM and FliN, form the C-ring, and are responsible for switching of the rotation between clockwise and counterclockwise (Morimoto and Minamino, 2014; Minamino and Imada, 2015).

The assembly of the flagellar motor has been mainly investigated in the model systems *E. coli* and *S. tiphymurium*. In these peritrichous bacteria, the initial component to form is the MS-ring, followed by the C-, P-and L-rings. A T3SS-like export apparatus is recruited by interaction with the MS-ring and is responsible for secretion of the single components of the rod, hook and filament, which are then assembled outside the cell (Minamino et al., 2008; Minamino and Imada, 2015; Nakamura and Minamino, 2019).

Whilst the MS-ring is recruited first to initiate flagellar biogenesis, it remains unclear which factors are needed for its recruitment and assembly. In *Salmonella typhimurium* it was observed that FliF overexpression leads to spontaneous assembly of MS-ring structures (Suzuki et al., 2004; Kawamoto and Namba, 2017; Kawamoto et al., 2020; Johnson et al., 2021) whereas in *Vibrio alginolyticus* the same behavior was not observed (Terashima et al., 2020). Furthermore, co-expression of FlhF and FliG promotes formation of MS-rings in *Vibrio* species (Terashima et al., 2020). These findings are in agreement with previous studies where it was highlighted that FlhF and FlhG are involved in regulation of flagellar localization and assembly in species with polar flagella and in some peritrichous species such as *Bacillus subtilis* (Kazmierczak and Hendrixson, 2013). FlhG is a MinD-like ATPase, and interacts with components of the C-ring, FliM, FliN and FliY (Schuhmacher et al., 2015a, 2015b). Upon interaction with these proteins, FlhG promotes their interaction and assembly with FliG (Schuhmacher et al., 2015a, 2015b). FlhF is a SRP-type GTPase that localizes at the cell pole to positively regulate the localization and formation of the flagellum by recruiting FliF (Green et al., 2009; Terashima et al., 2020), whereas FlhG was shown to act as a negative regulator of flagellar assembly through interaction with FlhF (Kusumoto et al., 2008; Kojima et al., 2020).

The structure of FliF in isolation was recently determined, and revealed that the RBM3 has a symmetry that can vary from C32 to C35, with the majority of particles displaying a C33 symmetry (Johnson et al., 2020). Astonishingly, this study showed that RBM2 forms rings with a 21-fold or 22-fold symmetry, localized at the inner part of the M-ring (Johnson et al., 2020), revealing a symmetry mis-match between the domains. RBM1 was not resolved in these structures. Subsequently it was shown that the prevalent symmetry for the basal body is C34 and that the RBM2 adopts preferentially a C23 symmetry at the internal face of the M-ring (Kawamoto et al., 2020). The cryoEM structure of the intact basal body has further confirmed that the RBM3 unambiguously displays a C34 symmetry (Kawamoto et al., 2020; Johnson et al., 2021). Nonetheless, these structures raised the question of how this protein can form an oligomeric assembly with different symmetries in different domains, and how their assembly is coordinated.

Here, we studied the oligomerization of the different FliF domains in isolation. We show that a construct encompassing RBM1, RMB2 and RBM3, but lacking the two trans-membrane helices, is still able to form the proper MS ring assembly, in the *Salmonella* orthologue (but not the *helicobacter* one). We demonstrate that the RBM2 and RBM3 domains oligomerize in isolation, and form ring-like structures, with symmetry corresponding to that of these domains within the basal body. In contrast, RBM1 in isolation is strictly monomeric. Intriguingly, we also report that a construct encompassing both RBM1 and RBM2 is monomeric, suggesting that within this construct, RBM1 prevents RBM2’s oligomerization. Finally, ectopic addition of RBM3 promotes the oligomerization of the RBM1-RBM2 construct, reversing the inhibition of RBM2 oligomerization by RBM1. Taken together, these results suggest that the oligomerization of FliF is coordinated and allow us to propose a model for the regulated formation of the MS ring to the final, correct assembly. This might play a role to prevent the premature formation and/or mislocalization of the flagellum basal body complex.

## 2 Results

### Oligomerization of individual domains of FliF

Previous studies have shown that when purified in isolation, the *S. typhimurium* FliF adopts its oligomeric state, including an unusual symmetry mis-match between RBM2 and RBM3 (Johnson et al., 2020; Kawamoto et al., 2020), suggesting a complex folding and assembly pathway for the MS ring. This observation prompted us to investigate if the individual RBMs could oligomerize on their own.

To this end, we engineered a series of constructs that encompass one or several RBMs (Figure 1b, Table 1). For each construct, the correspondent protein was purified, and its oligomerization propensity was analyzed by size exclusion chromatography (SEC) (Table 1, Figure 2a).

**Figure 2.**
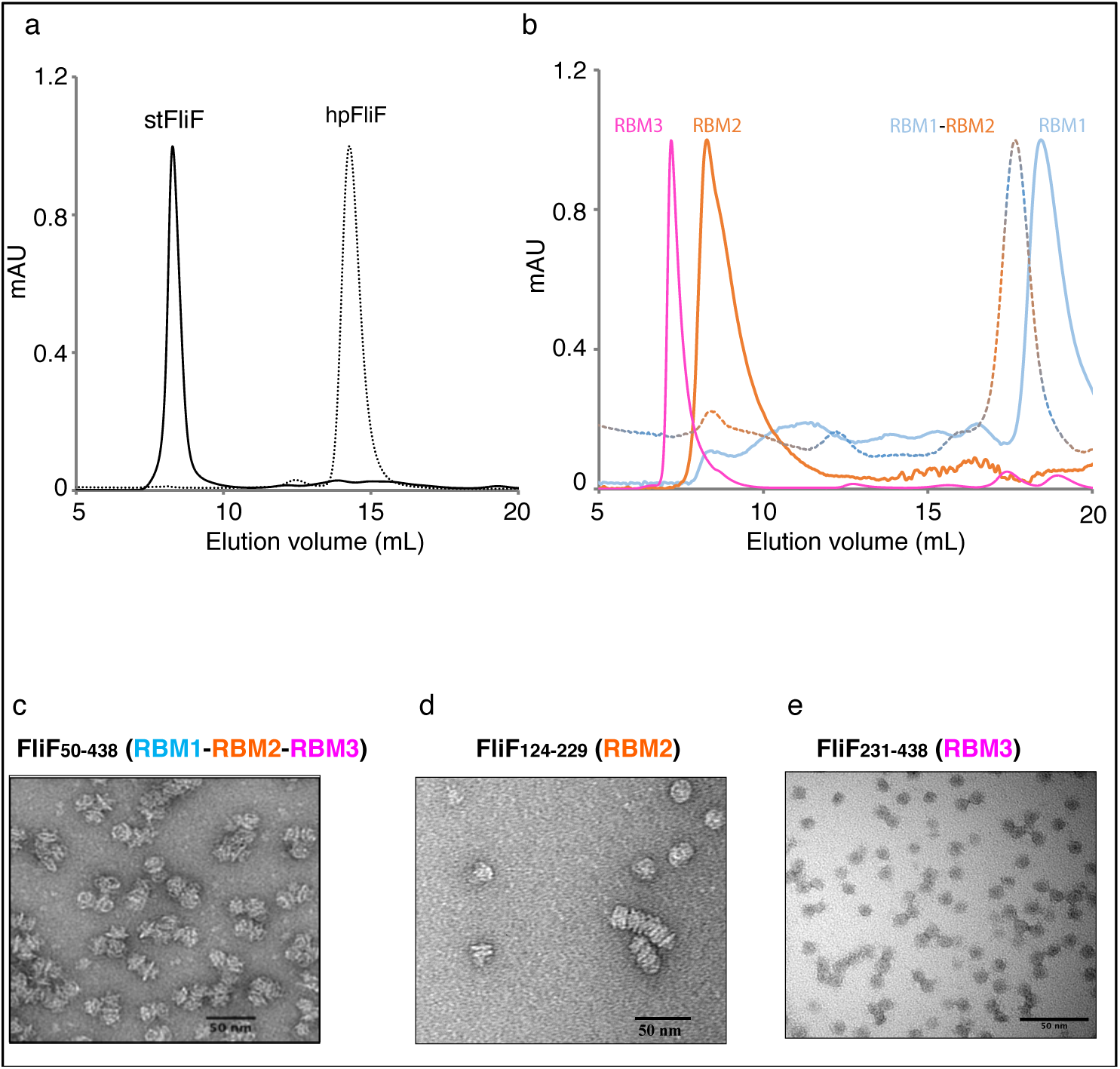
Oligomerization of the FliF domains. (a) Size exclusion chromatography UV trace of constructs encompassing the entire periplasmic regions of *S. typhimurium* FliF and *H. pylori* FliF. (b) Size exclusion chromatography UV trace of constructs encompassing the individual domains of *S. typhimurium* FliF (c-e) Negative stain analysis of (c) RBM1-RBM2-RBM3 (FliF_50-438_), (d) RBM2 (FliF_124-229_) and (e) RBM3 (FliF_231-438_). RBM1-RBM2-RBM3 and RBM2 shows mostly side views, whilst RBM3 mainly displays top views.

As shown on Figure 2a, a construct encompassing RBM1, RBM2 and RBM3 (FliF_50-438_) forms a high-order oligomer, stable by SEC. Negative-stain analysis revealed that the protein possesses ring-like features (Figure 2c), similar to that of the full-length protein. This demonstrates that the TM helices of FliF are dispensable for its oligomeric assembly. We however note that the protein is prone to aggregation, with multiple MS rings assembling from the side opposing the collar region, suggesting that some hydrophobic surfaces, possibly facing the membrane, are exposed in the absence of the TM helices. Indeed, SEC-MALS analysis confirmed that whilst FliF from *S. typhimurium* (StFliF_50-438_) self-oligomerizes in a complex with an apparent mass of ∼10 MDa (Figure S1a), significantly larger than the FliF 34-mer.

Next, we observed that constructs encompassing RBM2 (FliF_124-229_) or RBM3 (FliF_231-438_) also formed higher-order oligomers in isolation (Figure 2b). Negative-stain EM analysis confirmed that they form ring-like structures (Figure 2d, 2e), consistent with their architecture within the native MS ring. In the instance of RBM2 (FliF_124-229_), we note that the ring-like structures exhibited a tendency to cluster together, forming lines of discs (Figure 2d). It is noteworthy that in the T3SS FliF homologue SctJ, previous biochemical studies have shown that RBM2 is monomeric, and requires the L1 linker to oligomerize in isolation (Bergeron et al., 2015). This might suggest functional differences between the assembly of the T3SS and flagellum basal body.

In contrast to RBM2 (FliF_124-229_) and RBM3 (FliF_231-438_), we observed that the construct encompassing RBM1 (FliF_50-124_) is strictly monomeric in isolation (Figure 2b, Table 1). Collectively, these results demonstrate that in S. typhimurium, the TM helices of FliF are dispensable for its oligomeric assembly, and that RBM2 and RBM3, but not RBM1, can form oligomeric rings in isolation.

Whilst we observed that RBM1-RBM2-RBM3 (FliF_50-438_) in *S. typhimurium* spontaneously oligomerizes, previous studies have shown that in other non-peritrichous organisms, such as *Aquifex aeolicus*, RBM1-RBM2-RBM3 is strictly monomeric (Takekawa et al., 2021). For this reason, we investigated the oligomeric state of FliF in another non-peritrichous organism, *Helicobacter pylori*. We observed that *H. pylori* RBM1-RBM2-RBM3 (HpFliF_51-427_) elutes from the gel filtration column much later than StFliF_50-_438, consistent with a monomeric protein (Figure 2a). SEC-MALS analysis showed that the molecular weight of this purified protein is 41 kDa, in agreement with the predicted molecular weight of a single monomer. This result suggests that in non-peritrichous organisms, FliF might require additional factors to trigger oligomerization (Dasgupta et al., 2003; Hendrixson and DiRita, 2003).

### Cryo-EM analysis of the FliF RBM2 and RBM3

The structures of FliF revealed a range of stoichiometries, from 32 to 34 for RBM3, and 21 or 22 for RBM2, with an extra 11-12 RBM2 domains in a distinct orientation relative to RBM3, and facing outward (Johnson et al., 2020). Subsequent structures of this protein in the intact basal body demonstrated that the true stoichiometries are 34 and 23, respectively (Kawamoto et al., 2020; Johnson et al., 2021). This prompted us to use cryo-EM to characterize the oligomeric constructs described above, to confirm that they match the structure of the native FliF oligomer, and determine the stoichiometry of the individual domains.

As shown on Figure 3a, the FliF construct encompassing RBM1, RBM2 and RBM3 (FliF_50-438_) was readily incorporated into ice, which allowed us to collect a cryo-EM dataset. Because of the high level of aggregation (see above), we picked particles from this data manually, and used these to generate 2D classes (Figure 3b). These 2D classes are highly similar to that of the MS ring in isolation, with density for RBM2, RBM3 and the **β**-collar clearly visible (Figure 3c). Diffuse density below RBM2 is also visible, and was also seen in 2D classes of the MS ring, corresponding to density for dynamic RBM1 domains.

**Figure 3.**
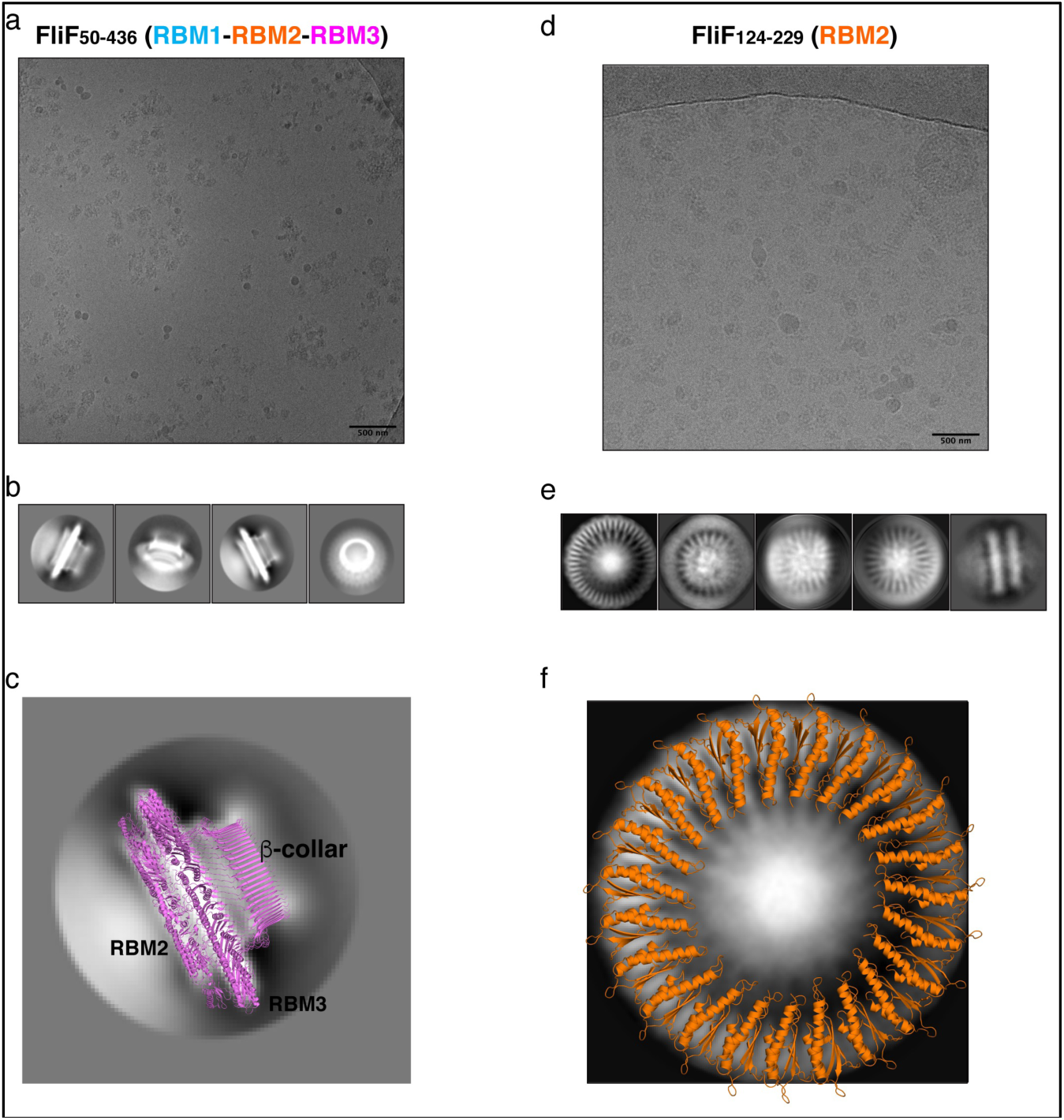
FliF RBM2 and RBM3 stoichiometry. (a) Cryo-electron micrograph and (b) selected 2D classes of RBM1-RBM2-RBM3 (FliF_50-438_). (c) A 2D class from a side view of RBM1-RBM2-RBM3 is shown, overlayed to a cartoon representation of the 33-mer FliF structure (PDBID: 6SD1). The majority of particles are side views, and match the architecture of intact FliF. (d) Cryo-electron micrograph, and (e) selected 2D classes of RBM2 (FliF_124-229_). Particles show top views and some filaments that consist of aggregation of single RBM2 discs. (f) A 2D class of a RBM2 top view is shown, overlayed to a cartoon model of the 23-mer RBM2 inner ring structure (from PDBID: 7BK0).

While most particles were attributed to 2D classes corresponding to side-views of the complex, a subset (∼10 %) correspond to top views (Figure 3b, far right). Notably, in this class, we were able to clearly identify a 33-fold symmetry (Figure S2a). This is in agreement with the structure of FliF in isolation, reported previously (Johnson et al., 2020), where RBM3 adopted a 33-mer stoichiometry in the majority of particles. Further work will be required to determine if our construct also adopts a range of stoichiometries.

Next, we used cryo-EM to characterize the RBM2 (FliF_124-229_) oligomer. This protein was also readily incorporated into ice (Figure 3D), and we were able to collect a cryo-EM dataset. We attempted automated particle picking using a range of tools, but only cryOLO(Wagner et al., 2019), was able to pick both side and top views, in particular as the side views consisted of long aggregation of discs (see above). Using these particles, we generated 2D classes in Relion (Scheres, 2012)(Figure 3e). These confirmed that this protein has a pathological level of preferred orientation, with most particles visible from the top of the ring, and very few tilted or side views, with the side views clustered together, as seen in negative stain (see above). This precluded high-resolution structure determination, but allowed us to exploit the top views to infer the symmetry of the particles.

In the intact FliF structure in isolation, RBM2 forms two rings: One inner ring with 21 subunits, and one outer ring with 9 subunits. As shown on Figure 3f we can observe on these 2D classes clear density for the 2 helices of RBM2, notably with a 23-fold symmetry (Figure S2b). Additional classification, using a larger top-view dataset, would be required if this sample is heterogeneous and includes a range of symmetries, as observed for the intact FliF. Nonetheless, this demonstrates that the oligomers obtained for our RBM2 construct (FliF_124-229_) correspond to the inner ring alone, and does not include the outer ring.

Finally, we note that in the RBM2 (FliF_124-229_) 2D classes, some density is visible in the centre of the ring, which cannot be interpreted with the current structures of FliF. We propose that this density likely corresponds to some undetermined chemical that was co-purified with the protein. Further work will be necessary to determine the nature of this additional density.

Collectively, these observations confirm that the FliF trans-membrane helices are not required for it to adopt its native MS-ring architecture. In addition, we show that the RBM2 of FliF adopts the 23-mer inner ring conformation in isolation.

### RBM1 prevents the oligomerization of RBM2, and this effect is counteracted by RBM3

Previous work on the T3SS FliF homologue SctJ had shown that RBM2 self-oligomerizes, similarly to FliF, but this oligomerization is repressed in the presence of RBM1 (Bergeron et al., 2015, 2018). We therefore sought to verify if the RBM1 of FliF played a similar role. To that end, we engineered FliF constructs that encompass both RBM1 and RBM2 (FliF_50-229_). As shown on Figure 2a, size exclusion chromatography analysis showed that the resulting proteins are strictly monomeric (Table 1). This suggests that RBM1 prevents RBM2 from oligomerizing on its own.

In order to determine how RBM1 could inhibit RBM2 domains to oligomerize, we first performed co-evolution analysis to determine amino-acid residues that are potentially involved into the interaction between RBM1 and RBM2, using RaptorX Complex Contact prediction server (Zeng et al., 2018). As shown in Figure S3a, several regions of the protein, largely corresponding to the β-strand regions, showed significant co-evolution scores. Next, we employed the HADDOCK docking server to model the interaction between the two domains, using these residues as restraints in the docking process. This led to a cluster of models with low energy score, and where the two domains have their β-sheet facing each other (Figure S3b), in agreement with the co-evolution analysis. Furthermore, overlay of this model onto the RBM2 23-mer structure has RBM1 in the position of an adjacent RBM2 molecule (Figure S3c), providing a potential explanation of how the intramolecular contacts between RBM1 and RBM2 sterically obstruct the RBM2 oligomerization. This is consistent with our observation that the RBM2 oligomerization is inhibited in the context of the RBM1-RBM2 construct.

This effect mentioned above was observed in the context on a RBM1-RBM2 construct. This led to the question of whether the addition of ectopic RBM1(FliF_50-124_) onto assembled RBM2(FliF_124-229_) rings leads to their dissociation. To verify this, we titrated purified RBM1 (FliF_50-124_) against oligomeric RBM2 (FliF_124-229_), and used ns-EM to investigate if the ectopic addition of RBM1 disrupts the RBM2 oligomers (see above). As shown on Figure S4, we observed no changes in the architecture or density of the RBM2 (FliF_124-229_) oligomers, even in large excess of RBM1 (FliF_50-124_). This observation demonstrates that once the RBM2 ring is formed, it can no longer be disrupted by RBM1, and suggests that in the context of the RBM1-RBM2 (FliF_50-229_) construct, RBM1 prevents RBM2 oligomerization by binding to the ring oligomerization interface.

Given that RBM1-RBM2 (FliF_50-229_) was shown to be strictly monomeric, whilst RBM1-RBM2-RBM3 (FliF_50-438_) assembles into the MS ring (Figure 2, Table 1), we further investigated whether addition to RBM3 (FliF_231-438_) would prompt RBM1-RBM2 (FliF_50-229_) to oligomerize. To this end, purified RBM1-RBM2 (FliF_50-229_) and RBM3 (FliF_231-438_) were mixed (Figure 4a), and ns-EM was employed to test the formation of the intact MS ring. Surprisingly, while we observed presence of ring-like structures formed by RBM3 (FliF_231-438_) alone, we also observed the presence of long tubular structures. These are distinct in appearance from the lines of discs observed for our RBM2 construct (see Figure 2c), but also to the RBM1-RBM2-RBM3 oligomers (See Figure 2b). We therefore propose that these tubular structures correspond to stacks of RBM1-RBM2 oligomers, induced and possibly capped by RBM3. This would however require to be experimentally verified. Nonetheless, this observation suggests that while RBM1-RBM2 exists as a monomer, addition of RBM3 is the determinant factor that pushes towards assembly of FliF into an oligomeric state.

**Figure 4.**
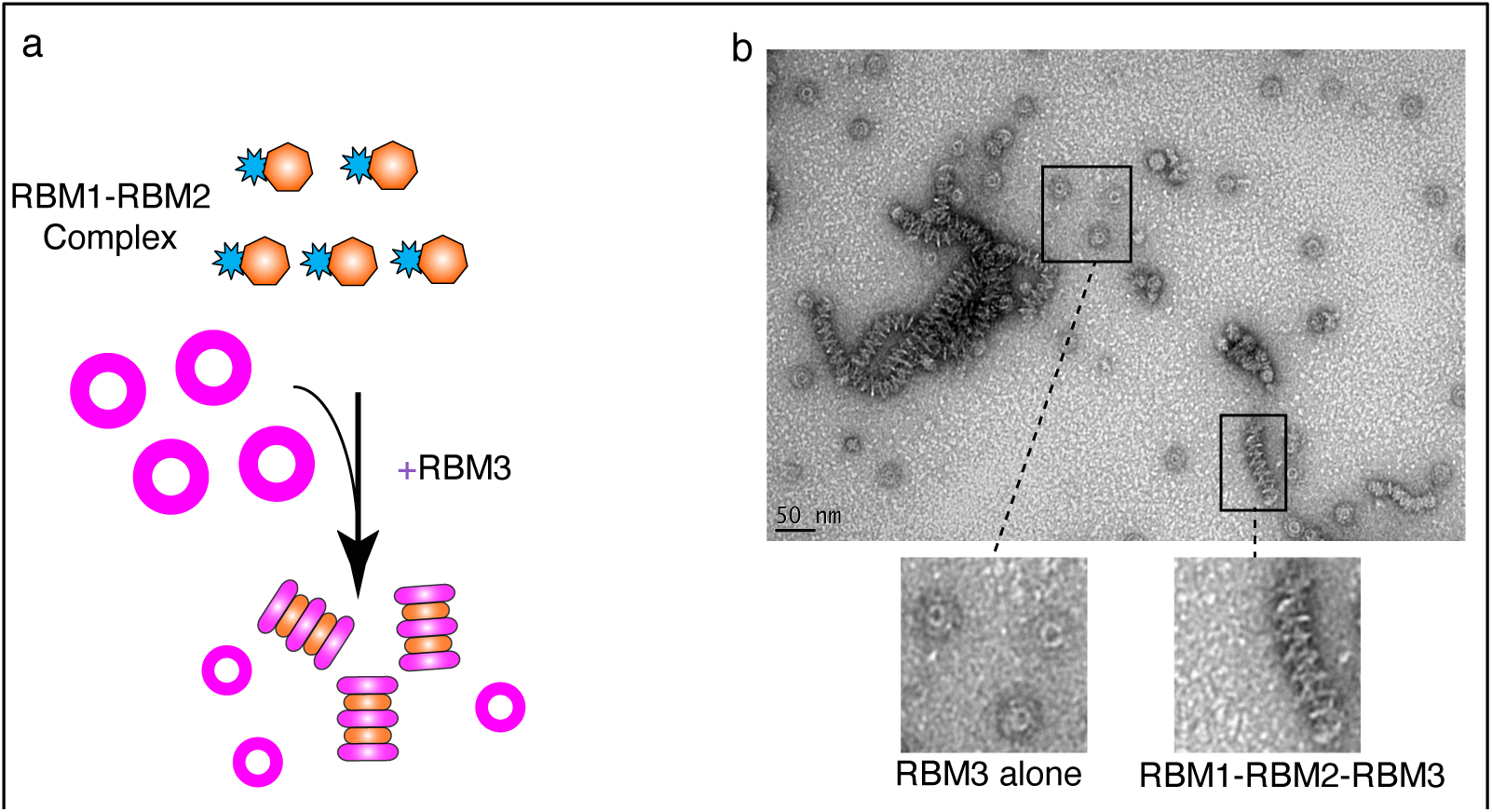
RBM3 induces oligomerization of the RBM1-RBM2 construct. (a) Schematic representation of the experiment. RBM1-RBM2 (FliF_50-229_) was mixed with RBM3 (FliF_231-438_) oligomers and imaged by negative stain EM. (b) Negative stain electron micrograph of the RBM3 and RBM1-RBM2 mixture. Ring-like structures, representative of RBM3 alone, are visible, as well as tubular structures that are likely composed of stacks RBM1-RBM2-RBM3 rings.

Collectively, these results suggest an intricate set of interactions between the different FliF domains, with RBM1 binding to RBM2 to prevents its oligomerization, and RBM3 acting to prevent this interaction.

## 3 Discussion

The MS-ring assembly is one of the first steps that occur during biogenesis of the flagellum (Minamino et al., 2008). The MS-ring then functions as a scaffold to recruit the C-ring through the interaction of FliF with FliG (Li and Sourjik, 2011; Morimoto et al., 2014) and the export apparatus (Minamino et al., 2008; Minamino and Imada, 2015; Nakamura and Minamino, 2019). Despite this central role, the process and regulation underlying the MS-ring folding remain unknown. A deeper understanding of FliF folding process has become increasingly important in light of the recent structural studies that have reported the existence of distinct symmetries within the MS-ring, which could serve multiple functions (Johnson et al., 2020, 2021; Kawamoto et al., 2020; Takekawa et al., 2021)

Indeed recent structural analyses have highlighted that the MS-ring symmetry can adopt a range of oligomeric states, with a mis-matched symmetry between RBM2 and RBM3 (Johnson et al., 2020). Whilst initially this had shown that RBM3 adopts a range of stoichiometries that range from 32 to 34 subunits, and that RBM2 forms either 21 or 22-mers (Johnson et al., 2020), in subsequent studies it was consistently observed that RBM3 is a 34-mer and RBM2 is a 23-mer (Kawamoto et al., 2020; Johnson et al., 2021). The symmetry mis-match between RBM2 and RBM3, together with the different symmetries detected in the existing studies suggests the existence of a complex process that regulates the folding and biogenesis of the MS-ring. In this study we aimed to determine the mechanism underlining the complex folding of FliF, by analysis the oligomeric state of the different domains of FliF.

Here we showed that in a construct encompassing FliF RBM1, RBM2 and RBM3 is able to assemble to form MS-rings, wherein RBM3 displays a 33-mer stoichiometry. Additionally, our data showed that RBM2 is able to form rings with a 23-mer stoichiometry. These correspond to the main stoichiometry observed for FliF in isolation. Conversely, we observed that a construct encompassing RBM1-RBM2-RBM3 (HpFliF_51-427_) from *H. pylori* yields a monomeric protein. These findings are in agreement with what was observed for FliF in *A. aeolicus* and suggest the existence of a different regulation of the MS-ring assembly for non-peritrichous organisms (Takekawa et al., 2021).

Indeed, our data demonstrates that constructs encompassing RBM1 and RBM2 are monomeric, conversely to what consisting of only RBM2. Since the addition of RBM1 to already formed RBM2 rings did not show any changes, we propose that RBM1 prevents the RMB2 oligomerization by binding to, and thus occluding, its oligomerization interface. Additionally, we also have shown that addition of RBM3 to monomeric RBM1-RBM2 caused formation of tubular structures, which we attributed to stacked rings formed by RBM1-RBM2-RBM3. This in turns suggests that RBM3 interacts with the RBM1-RBM2 construct in a way that dislodges RBM1, and allows RBM2 to oligomerize.

Based on this, we propose the following mechanism for MS ring assembly: Upon membrane insertion by the SEC pathway, FliF is a monomer; the interaction between RBM1 and the oligomerization interface on RBM2 retains FliF in a monomeric state (Figure 5a). Next, while RBM1 still prevents RBM2 molecules from associating, RBM3 oligomerization initiates (Figure 5b), imposing an overall 34-mer to the complex. Assembled RBM3 rings can subsequently disrupt RBM1 from RBM2 oligomerization interface, and RBM2 rings start forming (Figure 5c). These form 23-mer rings, but because the overall stoichiometry is imposed by the initial RBM3 oligomerization, 11 RBM2 domains are left on the outside. Therefore, we propose that the role of RBM1-mediated inhibition of RBM2 oligomerization in the FliF assembly process allows RBM3 rings to form and drive the MS-ring biogenesis process, determining the right stoichiometry for all the sub-assemblies. This leads to the formation of the intact MS-ring, with its symmetry mis-match between RBM2 and RBM3.

**Figure 5.**
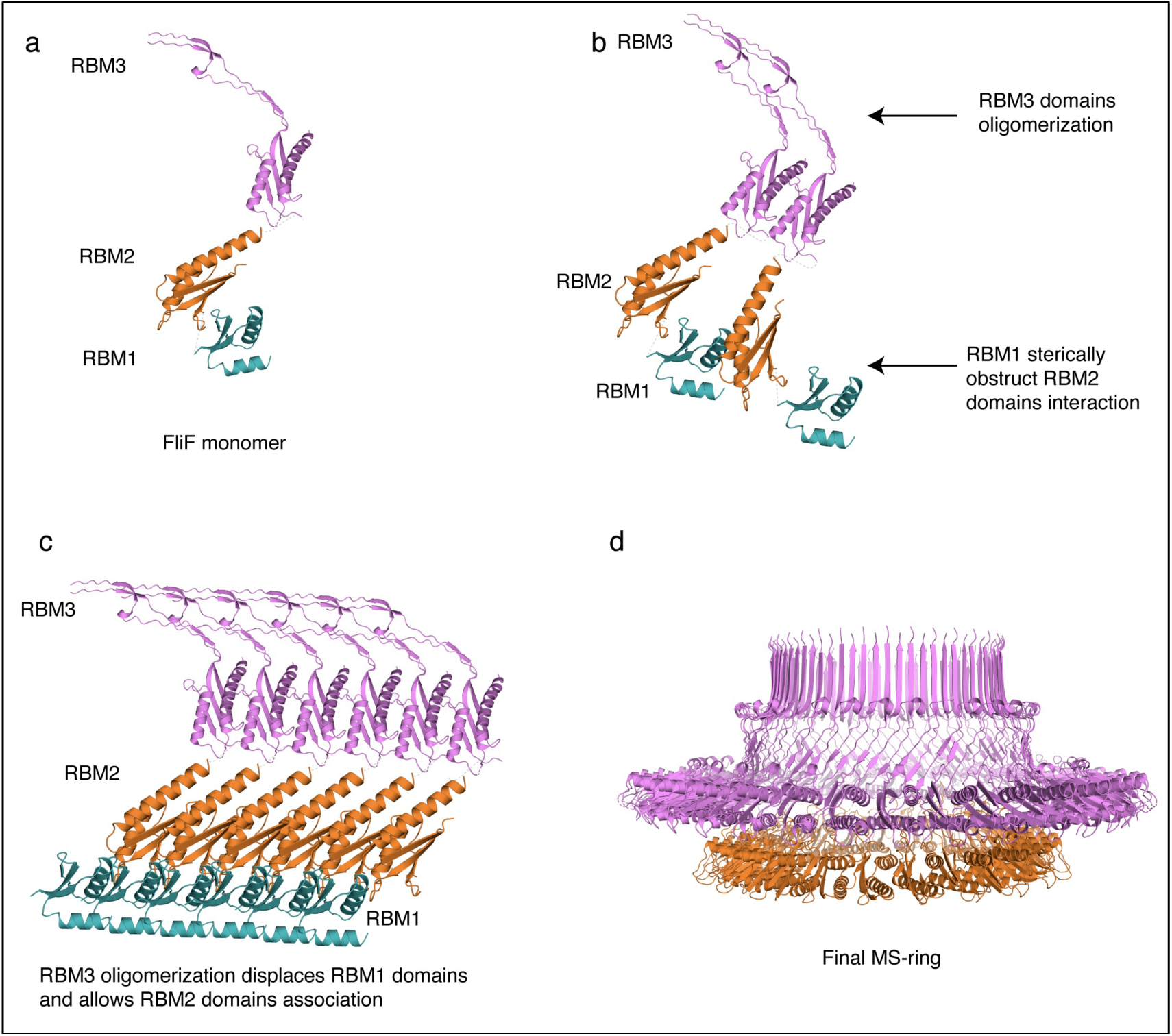
Structural representation of the proposed MS-ring assembly process. Cartoon representations of each domains, coloured as in Figure 1, are used. (a) As it gets exported to the membrane via the Sec pathway, FliF is monomeric, (b) The presence of adjacent FliF molecules allows RBM3 to drive the oligomerization process, establishing the 34-mer stoichiometry. The contacts between RBM1 and RBM2 prevent RBM2 to oligomerize. (c) At completion of RBM3 oligomerization, RBM1 is displaced from RBM2, allowing RBM2 domains from different FliF molecules to come into contact and assemble, forming a 23-mer. The remaining 11 RBM2 domains sit outside this ring. (d) In the final stage, the MS-ring reaches the final correct conformation, with a symmetry mismatch between RBM2 and RBM3.

The concept that the three periplasmic domains RBM1-RBM2-RBM3 of FliF might provide regulation of its oligomerization, thus guaranteeing the right stoichiometry of the MS-ring and the consequent correct assembly of the basal body is not foreign. Indeed a similar regulation has been proposed in the evolutionarily-related T3SS secretion apparatus (Yip et al., 2005; Bergeron et al., 2015, 2018; Bergeron, 2016). There, the proposed model suggests that the SctJ linker between RBM1 and RBM2 interacts with RBM1 with hydrophobic interaction, keeping SctJ in a monomeric state (Bergeron et al., 2015, 2018). Upon dissociation of the linker region from RBM1, SctJ subunits can associate establish a series of interactions between their respective RBM1-RBM2 domains, as well as the linker region state (Bergeron et al., 2015, 2018). SctD molecules subsequently insert between two adjacent SctJ subunits, and SctJ-SctD heterodimers can finally oligomerize to form the finalized rings (Bergeron et al., 2015, 2018). Our data show that whilst RBM1 and RBM3 can provide regulation of RBM2 oligomerization, it appears that the process is opposite to what observed in SctJ.

It is worth to note that while for SctJ the regulation role was pin-pointed to the linker region, in our case the FliF RBM1 region we used in our study encompassed both RBM1 and the linker between RBM1 and RBM2 and thus, further work will be needed to determine whether the inhibition of RBM2 oligomerization is determined by RBM1 or its linker. Nevertheless our data demonstrate that the initial RBM1-RBM2 interaction and the timely formation of RBM3 rings are fundamental steps that lead to the correct assembly of the MS-ring.

The biogenesis of the flagellum is a hierarchical process that initiated with the insertion of the Type III export apparatus and the assembly of the MS-ring. The remaining flagellar components are then secreted through the export apparatus to build up the final flagellar structure (Yonekura et al., 2002; Macnab, 2003). The levels of regulation of this process are complex, relying on the hierarchical and timely transcription of the distinct components of the flagellum, which are transcribed in different groups according to their role in the flagellar structure (Kutsukake et al., 1990; Dasgupta et al., 2003). In a similar fashion, it is possible to speculate that RBM1 and RBM3-mediated control over the oligomerization and assembly of the MS-ring will provide an additional level of complexity to the flagellum biogenesis.

Several studies have shown that the regulation process involves different factors between peritrichous and polar flagella. Namely, FlhF and FlhG are not present in *E. coli* and *S. tiphymurium* but are necessary for flagellar synthesis and localisation in a number of species (Pandza et al., 2000; Hendrixson and DiRita, 2003; Niehus et al., 2004; Kusumoto et al., 2008; Schuhmacher et al., 2015a). Interestingly, FlhF and FlhG were found to antagonistically influence the levels of expression of the distinct groups of genes involved in flagellum synthesis (Dasgupta et al., 2003; Hendrixson and DiRita, 2003). It is also noteworthy that in some species carrying FlhF and FlhG, FliF was found to remain in a monomeric state *in vitro* and oligomerization occurred only in presence of FlhF and FliG (Terashima et al., 2020). In this study we reported that FliF in *H*.*pylori* exists in a monomeric state *in vitro*, in accordance to what also observed for FliF in *A. aeolicus* (Takekawa et al., 2021). Given the non-peritrichous nature of the flagella of these two organisms, it is possible to speculate that they may also require FlhF and FlhG to trigger FliF oligomerization and it will require further investigation.

Conversely, in *S. tiphymurium* it has been shown that FliF can oligomerize spontaneously (Minamino et al., 2008; Johnson et al., 2020). These observations underline that different, multi-faceted mechanisms of regulations exist for correct assembly of the flagellar machinery between species and that control of FliF oligomerization in *S. tiphymurium*, provided by FliF own domains, adds a new level of complexity to the modulation of the flagellum biosynethesis. Ultimately, characterization of the differences in the assembly of the flagellum between species will provide a better understanding of the molecular elements that determine regulation of the flagellum

## 4 Material and Methods

### Protein expression and purification

The gene coding for *FliF* encompassing RBM1, RBM2 and RBM3 (FliF_50-438_) was synthesized (Bio Basic), and cloned into the pET-28a vector, to include with a Thrombin-cleavable N-terminal His_6_ tag. Other FliF constructs (see figure 1b) were generated by site-directed mutagenesis, using the aforementioned construct as a template.

For protein over-expression, the corresponding plasmids were transformed into *Escherichia coli* BL21 DE3 cells and grown at 25°C at 160 rpm overnight in ZYM-5052 auto-induction media (1% Tryptone, 0.5%Yeast Extract, 25 mM Na_2_HPO_4_, 25 mM KH_2_PO_4_, 50 mM NH_4_Cl, 5 mM Na_2_SO_4,_ 2 mM MgSO_4_, 0.5% glycerol, 0.05% glucose, 0.2% α-lactose) for 16h. Following induction, cells were centrifuged at 5000 x g and pellets resuspended in buffer A containing 50 mM Hepes pH 8.0, 500 mM NaCl, 20 mM imidazole. Cells were lysed by sonication following addition of cOmplete™ EDTA-free protease inhibitor (Sigma) and debris removed by centrifugation at 14,000 × g for 45 min. The cleared supernatant was applied onto a 5 ml ml HisPure™ Ni-NTA resin (Thermo Scientific) gravity-based column equilibrated with 10 column volumes of buffer A. Proteins were eluted with a 2 step-gradient elution containing 50 mM and 500 mM imidazole, respectively. Fractions containing purified FliF RBM2 (FliF_124-229_) were further purified by size exclusion chromatography using a Superdex 200 10/300 column (GE Healthcare) in a buffer containing 50 mM Hepes pH 9.0, 500 mM NaCl. Purified FliF RBM1-RBM2 (FliF_50-229_) and RBM3 (FliF_231-438_) were applied to a Superdex 200 10/300 column and to a Superose 6 10/300 column (GE Healthcare), respectively, in a buffer containing 50 mM Hepes pH 8.0, 500 mM NaCl.

### SEC-MALS analysis

Sample were run through a standard bore, 5 μ 300 Å SEC column (Wyatt), using an infinityII HPLC (Agilent), in buffer containing 20 mM HEPES pH 7.0, 150 mM NaCl, 1 mM DTT. MALS and DRI data were obtained uwing the DAWN and Optilab detectors, respectively (Wyatt), and analyzed with the Dynamics software (Wyatt) to determine the molecular mass.

### Negative-stain grid preparation and EM data acquisition

For negative-stain EM experiment, 5 μl of purified protein, at a concentration of 0.2 mg/mL were applied onto glow-discharged carbon-coated copper grids, and incubated at 20C for 2 min. The grids were then washed in deionized water, and incubated with 1% Uranyl Formate for 30 sec. For the titration experiment in Figure 4, FliF RBM1-RBM2 (FliF_50-229_) and RBM3(FliF_231-438_) were mixed at 1:1 ratio. For the titration experiment in Figure S2, RBM2 (FliF_124-229_) was kept at a constant concentration of 0.2 mg/mL, while RBM1(FliF_50-124_) was added at different ratios.

Images were acquired on a Technai T12 Spirit TEM (Thermo Fisher) equipped with an Orius SC-1000 camera (Gatan). For FliF RBM2 (FliF_124-229_) domain, images were acquired at a 49k magnification with a defocus range of −0.5 μm to −1.0 μm. For FliF RBM3 (FliF_231-438_) domain, images were acquired at a 30k magnification with a defocus range of −0.5 μm to −1.0 μm.

### Cryo-EM grid preparation, data collection and data processing

5μl of protein at a concentration of 10 mg/mL, in 50 mM Hepes (pH 9.0), 150 mM NaCl, was applied onto glow-discharged 300 mesh Quantifoil R1.2/1.3 grids. Grids were then blotted for 10 s at 80% humidity, and plunged into liquid ethane, using a Leica EM-GP plunge freezer.

For RBM2 (FliF_124-229_), micrographs were collected on a 300 kV Titan Krios microscope equipped with a Gatan K3 camera. 10053 movies were recorded with a pixel size of 0.85 Å with an exposure of 1 e^−^/ Å^2^/frame for 40-50 frames. For RBM1-RBM2-RBM3(FliF_50-438_), micrographs were collected on a 200 kV Tecnai Arctica equipped with a Falcon 3 camera. A total of 2540 movies were collected using a pixel size of 2.03 Å and an exposure of 0.8 e^−^/ Å^2^/frame over 50 frames.

Data processing was performed in RELION 3.1(Scheres, 2020). Motion correction was performed with MotionCor2 (Zheng et al., 2017). CTF parameters were estimated with CTFFIND4(Rohou and Grigorieff, 2015). For RBM2 (FliF_124-229_), 2000 micrographs were manually picked and used for training a model for particle picking in crYOLO 1.5 (Wagner et al., 2019). Trained model was then used for automated particle picking for the whole dataset and box files were imported on RELION 3.1 for particle extraction. A total of ∼ 2000000 particles were extracted with a 230 pixels box. Extracted particle were subjected to multiple rounds of 2D classification to filter top views that allowed evaluation of symmetry. For RBM1-RBM2-RBM3 (FliF_50-438_), automated picking was instead performed within RELION 3.1 and a total of 129.000 particles were extracted with a box size of 220 pixel.

### Sequence analysis and model docking

The co-evolution analysis between RBM1 (FliF_50-124_), and RBM2 (FliF_124-229_) was performed with the RaptorX Complex Contact prediction server (Zeng et al., 2018), using default parameters. To model the interaction between RBM1 (FliF_50-124_), and RBM2 (FliF_124-229_) based on the co-evolution data, we first generated a homology model of the *S. typhimurium* RBM1, based on the *A. aeolicus* RBM1-RBM2 crystal structure (PDB ID: 7CIK). We then employed the HADDOCK 2.4 server to predict the structure of a complex formed between this homology model and the RBM2 structure (from PDB ID 6SD4), with all the co-evolving residues with a score above 0.4 included as active residues in the interaction. 200 decoys were modelled, which could be classified in 10 clusters, three of which were very similar, with identical interaction interfaces and RMSD < 4A. These included the lowest-energy model, and combined represented 55 decoys, suggesting that it is likely close to the real complex structure.

## Acknowledgements

This work was funded by a UBC Centre for Blood Research Post-doctoral transition award, and by a UK Biotechnology and Biological Sciences Research Council (BBSRC) grant (BB/R009759/1), both to JRCB. Cryo-EM data was collected at the UK national Electron Bio-Imaging centre (eBIC), proposal EM19709-1, and at the University of Sheffield FoS Electron Microscopy Facility. We are grateful to Prof. Natalie Strynadka for use of her laboratory at the initial stage of the project, and to Prof. Susan Lea, Dr Emily Furlong and Dr Steven Johnson for useful discussion on the FliF symmetry. We thank John Hall (Wyatt) for assistance with the SEC-MALS data collection and analysis.

## Author contribution

RFR and JR cloned the various constructs; GM, RFR, SB, WZ, and JR purified the proteins; GM and RFR performed the negative-stain EM and cryo-EM analyses, and processed the cryo-EM data, with help from ST; JRCB conceptualized the project. GM and JB wrote the manuscript, with contributions from all authors.

