## Supplementary Figures for "Oligomerization of the FliF domains suggests a coordinated assembly of the bacterial flagellum MS ring"

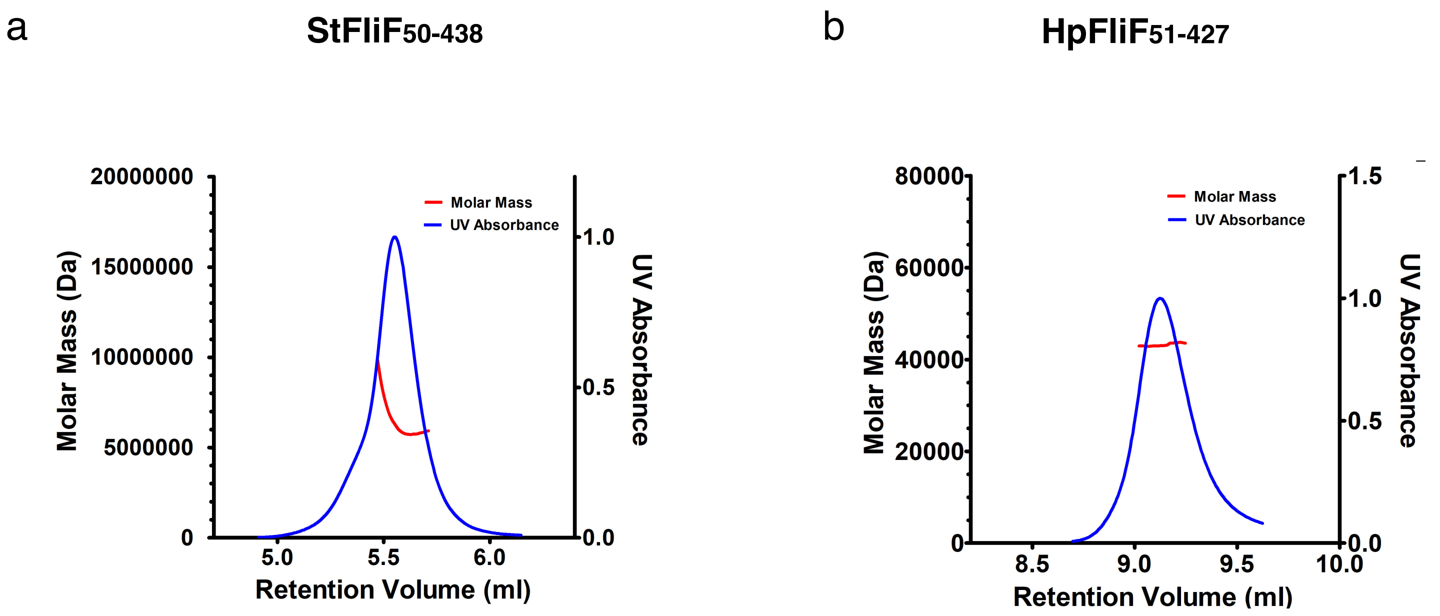


**Figure S1. SEC-MALS analysis of FliF from *S. typhimurium* and *H. pylori*** (a) SEC-MALS chromatogram of S. typhimurium FliF ($StFliF_{50-438}$). (b) SEC-MALS chromatogram of *H.pylori* FliF ($StFliF_{51-427}$). Blue line shows the UV traces while the red lines across the UV traces shows the MALS-derived apparent masses.

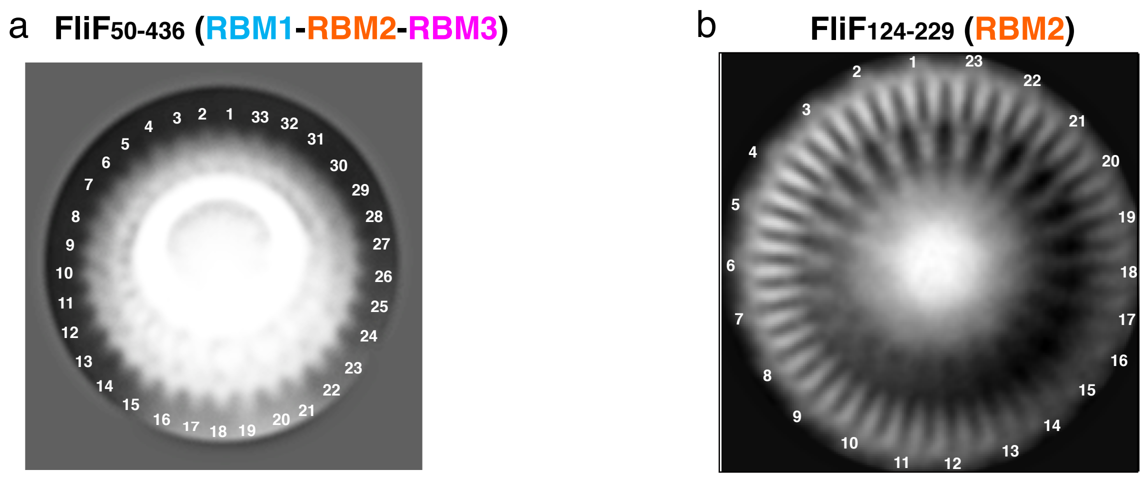


**Figure S2. Symmetry of RBM1-RBM2-RBM3 and RBM2 domains.** (a) Representative 2D class shows a top view of RBM1-RBM2-RBM3 (FliF$_{50\text{-}438}$) which displays C33 symmetry. (b) 2D class average of a RBM2 (FliF$_{124\text{-}229}$) top view shows C23 symmetry.

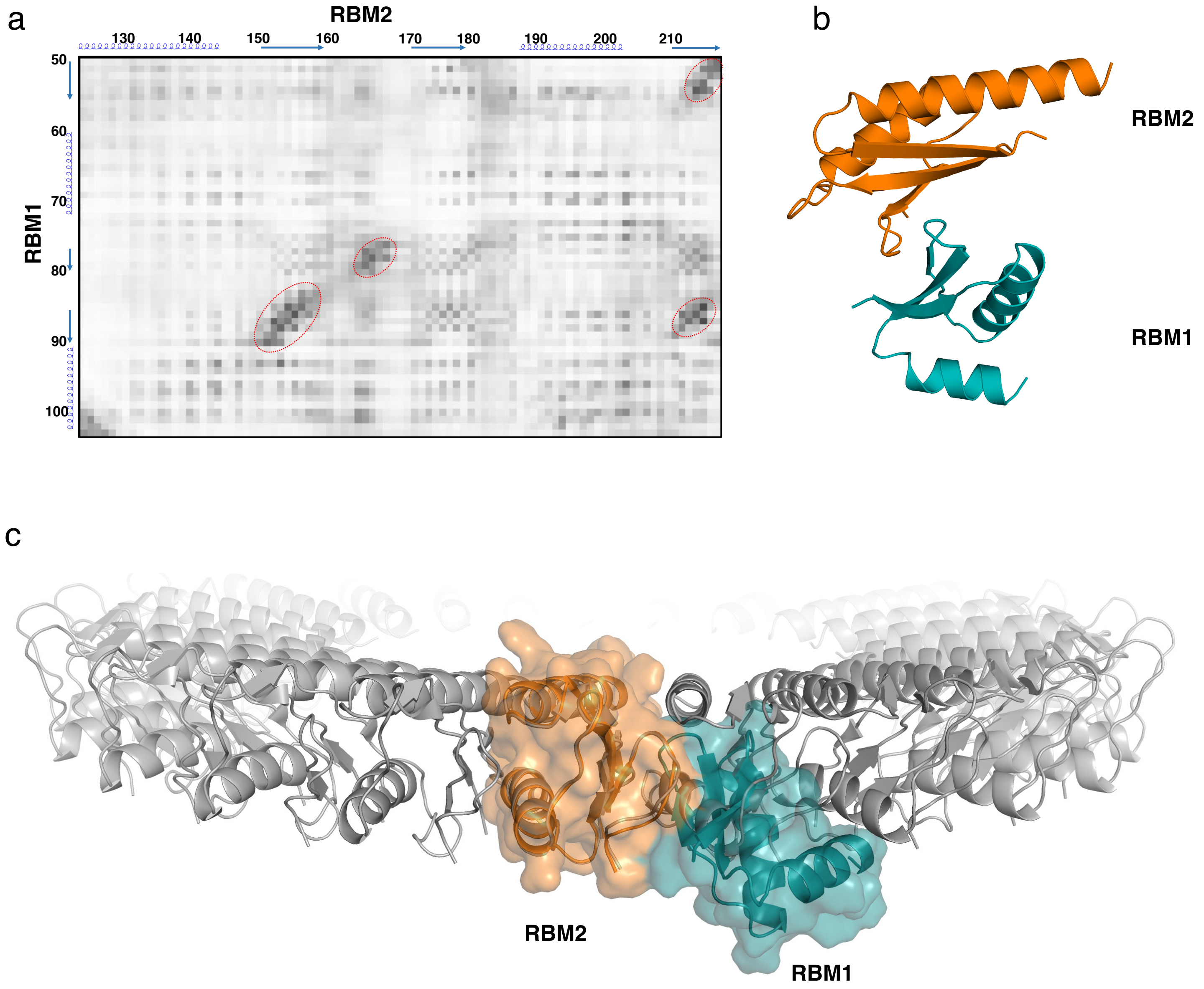


**Figure S3. Modeling of the RBM1-RBM2 interaction preventing RBM2 premature oligomerization.** (a) Heatmap of the predicted contacts between RBM1(FliF$_{50\text{-}124}$) and RBM2(FliF$_{124\text{-}229}$) obtained through co-evolution analysis on RaptorX software. RBM1 and RBM2 residues and secondary structure are shown next to the heatmap. Amino-acids likely involved in interactions between RBM1 and RBM2 are highlighted by red circles. (b) Cartoon representation of RBM1-RBM2 heterodimer predicted by HADDOCK, restrained using the residues with high co-evolution scores. Each domain is colored as in Figure 1. (c) The abovementioned RBM1-RBM2 model was overlayed to the RBM2 23-mer ring structure. The position or RBM1 sterically clashes with adjacent RBM2 molecules, consistent with the data showing that RBM1 prevents RBM2 oligomerization in the context of the RBM1-RBM2 construct.

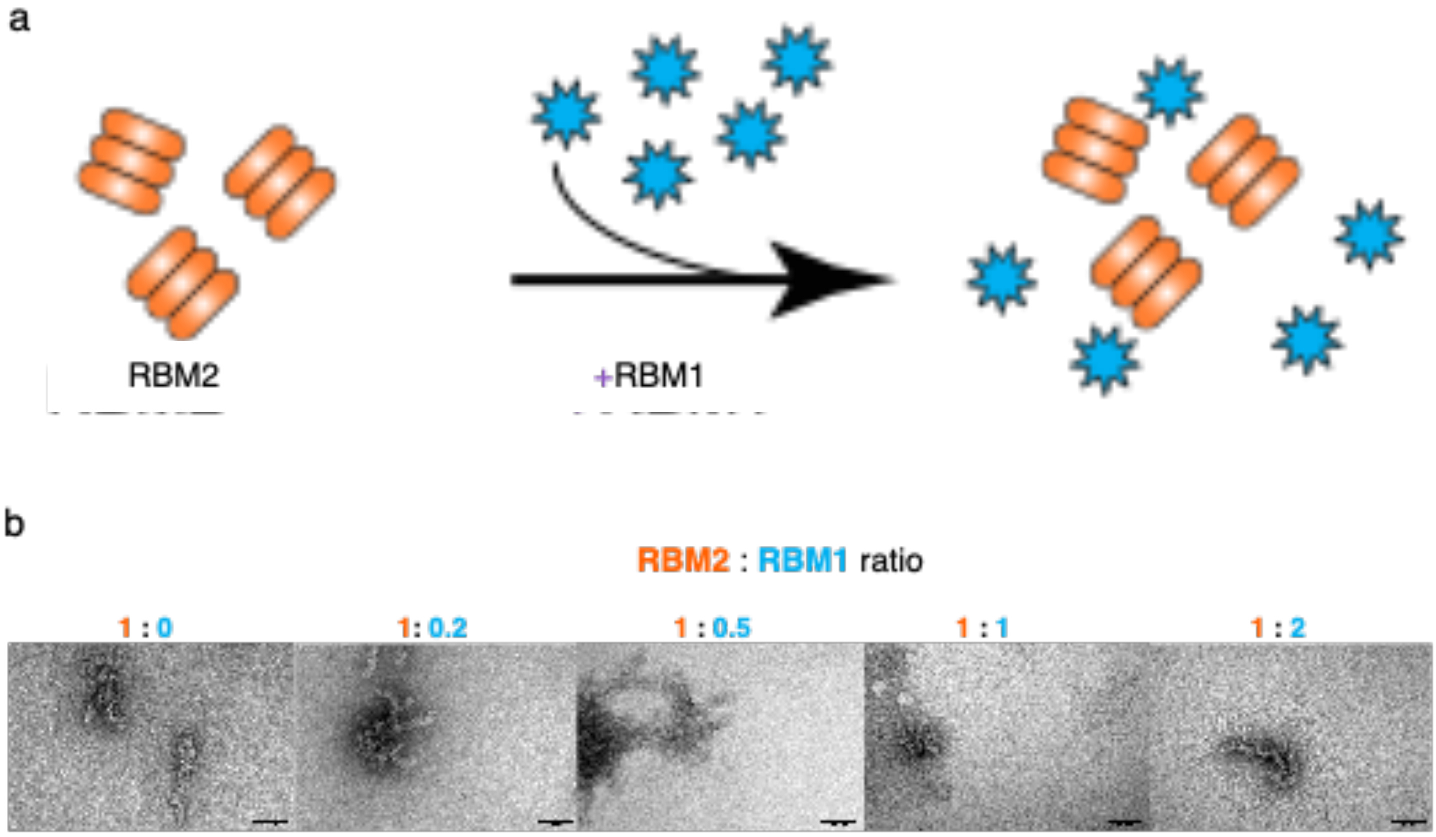


**Figure S4. Titration of RBM1 against RBM2.** (a) Schematic representation of the titration experiment. RBM2 ($FliF_{124-229}$) was mixed with increasing concentrations of RBM1 ($FliF_{50-124}$) and imaged by negative stain EM. (b) Negative stain electron micrographs of each point of the titration. Ratios at which RBM1 was added to RBM2 are shown on top of each micrograph. Addition of RBM1, even in large excess, does not disrupt RBM2 oligomerization.
