## Supplementary material for "Oligomerization of the FliF domains suggests a coordinated assembly of the bacterial flagellum MS ring": Table 1

| Domain | Construct boundaries | Monomer MW (KDa) | Oligomeric state of the purified protein |
| --- | --- | --- | --- |
| RBM1+L1 | 50-124 | 10.98 | Monomer |
| RBM2+L2 | 124-229 | 14.06 | Oligomer |
| RBM1+L1+RBM2+L2 | 50-229 | 22.09 | Monomer |
| RBM3 | 231-438 | 25.34 | Oligomer |
| RBM1+RBM2+RBM3 | 50-438 | 46.68 | Oligomer |
| Full length | 1-560 | 63.95 | Oligomer^[1]^ |

1. Johnson, Steven, Emily J. Furlong, Justin C. Deme, Ashley L. Nord, Joseph J. E. Caesar, Fabienne F. V. Chevance, Richard M. Berry, Kelly T. Hughes, and Susan M. Lea.‘Molecular Structure of the Intact Bacterial Flagellar Basal Body’. *Nature Microbiology* 6, no. 6 (June 2021): 712–21. https://doi.org/10.1038/s41564-021-00895-y.
